# MLKL D144K mutation activates the necroptotic activity of the N-terminal MLKL domain

**DOI:** 10.1101/2021.07.08.451689

**Authors:** Katja Hrovat-Schaale, Maruša Kalič-Prolinšek, San Hadži, Jurij Lah, Gregor Gunčar

## Abstract

Mixed-lineage kinase domain-like protein (MLKL) is an essential effector protein of necroptotic cell death. The four-helix bundle domain (4HB) presented by the first 125 amino acids of the N-terminal domain is sufficient for its necroptotic activity. However, it has been proposed that the subsequent helix H6 of the brace region has a regulatory effect on its necroptotic activity. How the brace region restrains the necroptotic activity of the N-terminal domain of MLKL is currently unknown. Here, we demonstrate the importance of helix H6 to constrain the necroptotic activity. A single amino acid mutation D144K was able to activate the necroptotic activity of the N-terminal domain of MLKL by removing helix H6 away from 4HB domain. This enabled protein’s oligomerization and membrane translocation. Moreover, a biophysical comparison revealed that helix H6 becomes partially unstructured due to D144K mutation, leading to a lower overall thermodynamic stability of the mutant protein compared to the wild type.

## Introduction

Necroptosis is a caspase-independent regulated type of cell death that is induced by specific inflammatory stimuli including tumor necrosis factor (TNFα)[1], death receptor [2], Toll-like receptor and interferon receptor agonists [3]. Due to its lytic nature, resulting in the release of intracellular damage-associated molecular patterns (DAMPs) [4], it is implicated in several inflammatory diseases of the skin [5, 6], eye [7], liver [6, 8], kidney [9], brain and intestinal tract [6]. The immunological role of necroptosis in response to pathogens remains unclear. Evidence suggest that it could act as a defense mechanism against viral infection when apoptosis is compromised [10]. This has been suggested in the case of vaccinia virus [1] and murine cytomegalovirus infections [11]. Contrary to this theory, certain intracellular bacteria, such as *Salmonella* Typhimurium [12] use the lytic nature of necroptosis to evade the immune system, suggesting an ambivalent role of the necroptotic pathway in host defense.

Receptor interacting protein kinase 1 (RIP1) [8, 13] and 3 (RIP3) [1, 14, 15] were identified as key regulators of necroptosis along with the pseudokinase mixed lineage kinase-domain like protein (MLKL) [16–19]. Even though RIP1 can be dispensable for necroptosis or can even inhibit necroptosis during a developmental process [6], there is no current evidence to suggest necroptosis could occur independently of RIP3 or MLKL [20].

MLKL contains N-terminal four-helix bundle domain (4HB), followed by a two α-helices termed the brace region and the regulatory C-terminal pseudokinase domain [19]. Phosphorylated RIP3 activates human MLKL by phosphorylating Thr357 and Ser358 within the C-terminal domain [16]. This results in a conformational change of MLKL, exposing its N-terminal domain. Overexpression of human 4HB domain alone (first 125 amino acid residues) in HEK293T and mouse 4HB (first 125 amino acid residues) domain in mouse dermal fibroblasts alone is sufficient to kill cells. Thus, 4HB domain is considered to be the executioner part of MLKL [21, 22].

The exact mechanism of necroptotic activity of the MLKL is still unknown. Several models have been proposed so far including activation of ion channels [23, 24], direct plasma membrane permeabilization and pore formation [22, 25–28]. Once N-terminal domain is exposed, MLKL forms oligomers. Trimers [21, 23, 29], tetramers [24, 30] hexamers [25], octamers [31] and even high molecular weight complexes [21, 22, 29, 32], have been reported. Even though the stoichiometry of the complex remains a matter of ongoing research, it is well accepted that MLKL forms oligomers and is translocated from cytoplasm to the plasma membrane where it disrupts membrane integrity. The N-terminal domain responsible for oligomerization [25, 27, 29], is also responsible for the binding of negatively charged membrane lipids including phosphatidylinositol phosphate (PIP) and cardiolipin [22, 25–27, 30]. Whether oligomerization and lipid binding are sufficient for MLKL membrane translocation or other factors are involved in this step remains to be discovered. Although the role of 4HB domain and C-terminal pseudokinase domain in MLKL’s activity have been relatively well investigated, little is known about the role of the brace region connecting both domains. Recently, the second brace helix of the N-terminal domain has been shown to play a role in MLKL’s oligomerization [23, 29]. Interestingly, overexpression of MLKL construct encompassing the 4HB domain and the first brace helix (residues 1-154), and liposome leakage assay of the similar construct (residues 2-154) indicated a possible inhibitory effect of the first brace helix on 4HB domain [26, 33]. However, the exact mechanism of how helix H6 could regulate the activity of MLKL has yet not been elucidated.

To gain insight into how helix H6 could regulate the necroptotic activity of the 4HB domain, we introduced a single point mutation that would repeal the helix H6. Aspartate 144 was identified as the most sensible candidate by sequence alignment and crystal structure inspection. First, we introduced a single amino acid mutation D144K into the inactive N-terminal domain of MLKL (MLKL_154_) and confirmed that this mutation did not disrupt protein’s secondary structure. Next, we showed that D144K mutation is sufficient to restore the necroptotic activity of the inactive MLKL_154_ in human HEK293T as well as in bacterial cells. Furthermore, we demonstrated that D144K mutation leads to a partial unwinding of the helix H6 and its displacement from the 4HB domain resulting in an overall decrease in protein thermodynamic stability compared to MLKL_154_. Finally, we showed that D144K mutation enables MLKL_154_ translocation to the cytoplasmic membrane and possibly its oligomerization. Our results support the hypothesis of a plug release mechanism [26] where 4HB is kept inactive by the first α-helix, helix H6 of the brace region [21, 22, 25, 27, 33].

## Materials and Methods

### Construct preparation

Full-length MLKL isoform 1 (MLKL1), the N-terminal domain MLKL_154_ and MLKL_139_ were cloned into pcDNA3.1 vector (Life Technologies) as previously described [33]. A single point mutation D144K was introduced into pcDNA3.1 vector coding for MLKL1 and MLKL_154_ using PCR site-specific mutagenesis. The DNA template for wild type and mutant MLKL_154_, as well as MLKL_139_, were sub-cloned into pMCSG7 vector with ligation independent cloning [34]. The DNA template for the mutant and the wild type MLKL_154_ were sub-cloned into pEGFPN2 (Life Technologies) vector using XhoI and BamHI (NEB) restriction sites. All constructs were sequence verified. Plasmid DNA was purified using Endofree Maxi kits (Qiagen) for use in cell-based assays. Primer sequences used for cloning are available upon request.

### Bacterial growth conditions and viability assay

The expression strain of *E. coli* BL21 [DE3] pLysS was transformed with pMCSG7 vector coding for MLKL_154_, MLKL_154_ D144K or anti-MLKL nanobody fused with mCherry, used as a negative control. The transformed cultures were grown in LB medium supplemented with ampicillin (100μg/ml) and chloramphenicol (50μg/ml) at 37°C and 130rpm to an optical density of 0.55. At time zero, protein expression was induced by addition of IPTG to a final concentration of 0.5mM and cells were grown for an additional 2h. Non-induced, transformed bacterial cultures were grown under the same conditions and were used as a control. Serial dilutions were prepared in the above mentioned LB medium at appropriate time points (0 and 2h) and were plated onto LB agar plates with appropriate antibiotics and grown overnight at 37°C for viable CFU counts. The relative percentage of bacterial viability was calculated according to the formula % of relative viability = [(CFU/ml of interest 2h post-induction)/(CFU/ml of CTRL two hours post-induction)]×100. Protein expression was confirmed 2h post-induction by immunoblotting.

### Mammalian cell culture and viability assay

HEK293T cells were cultured in DMEM medium supplemented with 10% FBS (Gibco), penicillin (50 units/ml) and streptomycin (50μg/ml) in a humidified atmosphere of 5% CO_2_ and 37°C. One day prior to transfection HEK293T were seeded at 2×10^4^ cells per well into 96-well plates. 100ng of a plasmid coding for a gene of interest was transfected using PEI transfection reagent (Sigma). Supernatants were collected 24h post-transfection, centrifuged for 5min at 500×g and LDH release was measured using the *In Vitro* Toxicology Assay Kit (Sigma-Aldrich) according to the manufacturer protocol. Total cellular LDH was determined by lysis of HEK293T cells with 0.1% TritonX-100. The absorbance at 490nm was measured using Powerwave XS microplate reader, and cell viability was calculated according to the formula % of cell death = [(released LDH – spontaneous LDH)/(total LDH – spontaneous LDH)]×100.

### Confocal microscopy

5×10^4^ HEK293T were seeded on poly-L-lysine (Invitrogen) coated coverslips (Roth) and cultured overnight. Cells were then transiently transfected with 250ng of empty pEGFPN2 plasmid or pEGFPN2 plasmids coding for MLKL_154_ or MLKL_154_ D144K and 250ng of empty pcDNA3.1 plasmid using PEI transfection reagent (Sigma). 24h post-transfection cells were fixed in 4% formaldehyde solution (Thermo Scientific) and mounted in antifade reagent with DAPI (Life technologies). Slides were analyzed at room temperature with Leica TCS SP8 confocal microscope under 63× oil immersion objective (NA=1.4). Excitation (488nm) and emission (510nm) wavelengths were used to visualize EGFP fluorophore and excitation (405nm), and emission (460nm) were used to visualize DAPI.

### Protein expression and purification

The recombinant proteins were expressed in *E. coli* BL21 [DE3] pLysS. Cells harbouring the expression plasmid coding for MLKL_154_ and MLKL_139_ were grown at 37°C in LB medium with ampicillin (100μg/ml) and chloramphenicol (50μg/ml) to an optical density of approximately 0.6. Protein expression was induced with IPTG at a final concentration of 0.5mM, and cells were grown under the same conditions for an additional 3 h. The recombinant MLKL_154_ D144K was expressed by an auto-induction medium as previously described (37). Briefly, overnight culture was prepared in a minimal MDG medium with ampicillin (100μg/ml) and chloramphenicol (50μg/ml), then diluted 1000-folds into auto-induction ZYM-5052 medium with ampicillin (100μg/ml) and chloramphenicol (50μg/ml) and grown at 18°C until the optical density reached a plateau (approximately 46h). Cells were then harvested, pelleted in binding buffer (20mM imidazole, 500mM NaCl and 50mM Tris-HCl pH 7.4) and stored at −80°C. Resuspended pellets were lysed by sonification, and soluble material was loaded onto an IMAC column (Ni-NTA agarose, GE Healthcare) pre-equilibrated with binding buffer. Unbound material was washed with binding buffer, and His_6_ tagged protein was eluted with elution buffer (300mM imidazole, 500mM NaCl and 50mM Tris-HCl pH 7.4). The N-terminal His_6_ tag was removed using TEV protease during overnight dialysis against cleavage buffer (50mM Tris-HCl pH 7.4, 150mM NaCl, 2mM β-mercaptoethanol). Cleaved His_6_ tag, uncleaved recombinant protein and TEV protease were removed by additional IMAC step. In the case of MLKL_154_ D144K, protein was diluted once with binding buffer before the final size exclusion purification step. The recombinant proteins were further purified by size exclusion chromatography in 150mM NaCl, 1.5mM DTT, 50mM Tris-HCl pH 7.4 buffer supplemented with complete EDTA-free protease inhibitor cocktail (Roche) using Superdex 75 10/300 column (GE Healthcare). The purity of the final samples was verified by SDS-PAGE analysis. For further analysis, MLKL_154_ was dialyzed against either 50mM Tris-HCl pH 7.4, or 20mM acetate buffer pH 5.0 containing 150mM NaCl, whereas MLKL_154_ D144 was dialyzed against the same buffers with 350mM NaCl.

### Size exclusion chromatography

To determine the oligomeric state of proteins, size exclusion chromatography was used. Purified recombinant proteins were loaded onto Superdex 75 10/300 column (GE Healthcare) in 150mM NaCl, 5mM Tris-HCl pH 7.4 and 1.5mM DTT.

### SDS-PAGE and immunoblotting

For analyses of protein expression in bacterial viability assay, total cell lysate loading was normalized based on the OD_600_ of cell culture at the appropriate time point. Briefly, OD_600_ was measured, and the sample concentration factor was calculated. An appropriate amount of bacterial suspension was pelleted by centrifugation at 10.000×g, 4°C for 4min. Pellets were resuspended in the appropriate amount of lysis buffer and lysed by sonification. 15μl of cell lysate was mixed with 5μl of 4× loading buffer, boiled for 3min and subjected to electrophoretic separation on denaturing polyacrylamide gels under reducing conditions, followed by transfer to nitrocellulose membranes. The latter were then probed with mouse anti-His HRP conjugated antibody (1:400, Sigma-Aldrich) and visualized by chromogenic DAB reagent (Sigma).

### Circular dichroism spectroscopy

Far-UV CD measurements were carried out on a JASCO J-1500 CD spectrophotometer (Jasco, Japan). All data were measured in 50mM Tris-HCl buffer pH 7.4 with 150mM NaCl for MLKL_154_ and 350mM NaCl for MLKL_154_ D144K and 1mM DTT with or without 30% (v/v) TFE. Protein samples were prepared either at 0.01mg/ml concentration and measured in quartz cells with 4mm path length or at 0.2mg/ml and measured in 1mm cuvettes. Total CD-spectra were measured at 25°C in the wavelength range of 200–250nm. Averaged spectra were calculated from three separate scans and measured with a step size of 1nm and a signal-averaging time of 4s. CD-signals were converted to mean molar ellipticity per residue [θ] using the following equation (1):

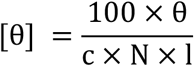

Where c is protein concentration (M), N is the amount of the amino acid residues and l is path length (cm). Secondary structure content was estimated using BeStSEL [35],[36].

### Intrinsic tryptophan fluorescence

Intrinsic fluorescence spectra were recorded in 10×4 mm quartz cuvettes at 25°C on a PerkinElmer LS 55 fluorimeter. All data were measured in 50mM Tris-HCl buffer pH 7.4 with 1mM DTT. Samples were excited at 280nm at a protein concentration 0.005mg/ml. Emitted fluorescence spectra were recorded from 300 to 420nm.

### Thermal denaturation and protein stability

Thermal denaturation profiles were obtained by recording the ellipticity at 222nm as a function of temperature in the range 4–94°C every 2°C with averaging time 4s. Protein samples were initially measured at pH 7.4 (50mM Tris-HCl, 150 or 350mM NaCl and 1mM DTT). However, reversibility was only partially restored under this condition. Reversibility was improved when protein samples were measured at pH 5.0 (20mM acetate, 150 or 350mM NaCl and 1mM DTT). MLKL_154_ was measured at 0.2mg/ml in 1mm cuvettes, and MLKL_154_ D144K was measured at 0.01mg/ml in 4mm cuvette. Thermal denaturation data measured at pH 5.0 were analyzed assuming a two-state model using a model equation that accounts for pre- and post-transitional baselines [37]. Fitting parameters: melting temperature at midpoint transition (*T_m_*), transition enthalpy (Δ*H_T_m__*) and heat capacity change (Δ*c_p_*), which was likely temperature-independent in the studies interval, were determined using nonlinear regression and extrapolated to 37°C according to Gibbs-Helmholtz relation. Since the curvature of melting transition did not permit reliable determination of Δ*c_p_* independently for MLKL_154_ D144K, we speculated that Δ*c_p_* increment is the same as for wild type N-termina domain of MLKL. Such an assumption is generally justified for single point mutations [38].

### Statistical analysis

Combined data from multiple independent experiments and containing at least three variables were subjected to ANOVA analysis, followed by Dunnett’s Multiple Comparison test.

## Results

### Amino acid residue at position 144 in helix H6 is well preserved among different species

Our goal was to pinpoint the amino acid most suitable to disrupt the interaction between 4HB domain and helix H6 of the brace region. We first searched for conserved amino acid residues on helix H6 among different species (Fig. 1B). There are 23 amino acids composing helix H6 in human MLKL [26]. Several of these amino acids are well conserved among 16 compared species, however only D140 and D144 were conserved among all compared species and were oriented towards 4HB. By sequence alignment and crystal structure inspection, we established that the most sensible candidate would be D144 (Fig. 1A). D144 is within the hydrogen bond distance to C24, and it also sits on the top of the helix H2. Helix H2 dipole and its two lysine residues (K25 and K26) strengthen this interaction. Therefore, we prepared a single amino acid mutation D144K.

**Figure 1:**
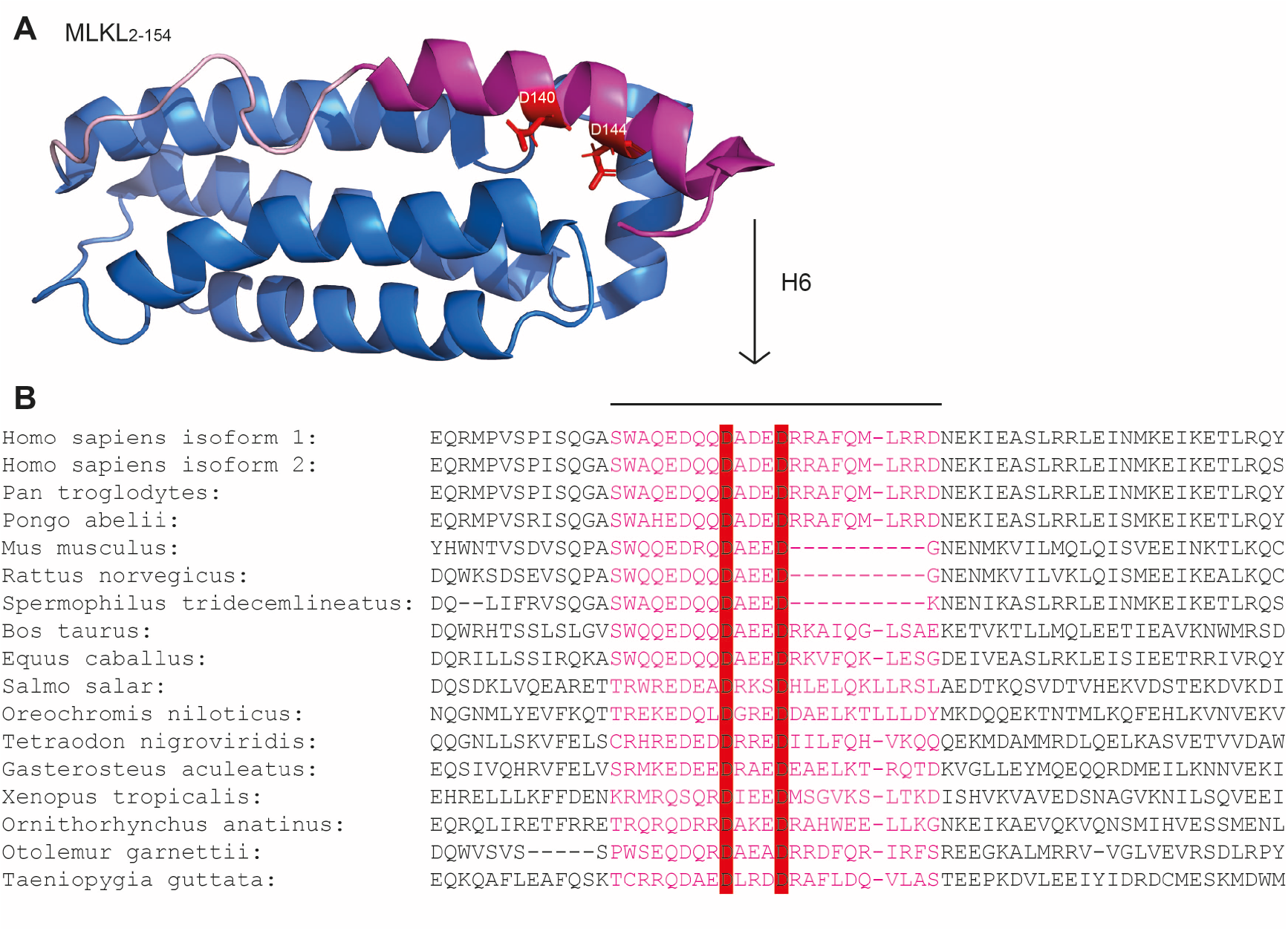
Amino acid residues D140 and D144 are conserved in helix H6 of MLKL among different species. (A) The structure of the N-terminal MLKL domain of MLKL_2-154_ shown in cartoon (PDB: 2MSV). 4HB is shown in blue, and H6 is shown in pink. (B) Amino acid residues alignment of the first α-helix of the brace region H6 shown in pink. D140 and D144 are highlighted in red.

### D144K activates the killing potential of the MLKL_154_ in bacterial and mammalian cells

Surprisingly, the growth of bacteria expressing MLKL_154_ D144K did not reach the optical density comparable to the bacteria expressing MLKL_154_. We therefore, performed bacteria viability assays as previously described [39] to see if there is a toxic effect of MLKL_154_ D144K on bacterial survival. Intriguingly, the mutant D144K lowed bacterial viability significantly compared to wild type MLKL_154_ 2h after protein induction. No such effect was seen on bacteria where protein expression was not induced (Fig. 2A, C). Since all expressed proteins were fused with His_6_ tag, their expression was confirmed with anti-His immunoblotting (Fig. 2B).

**Figure 2:**
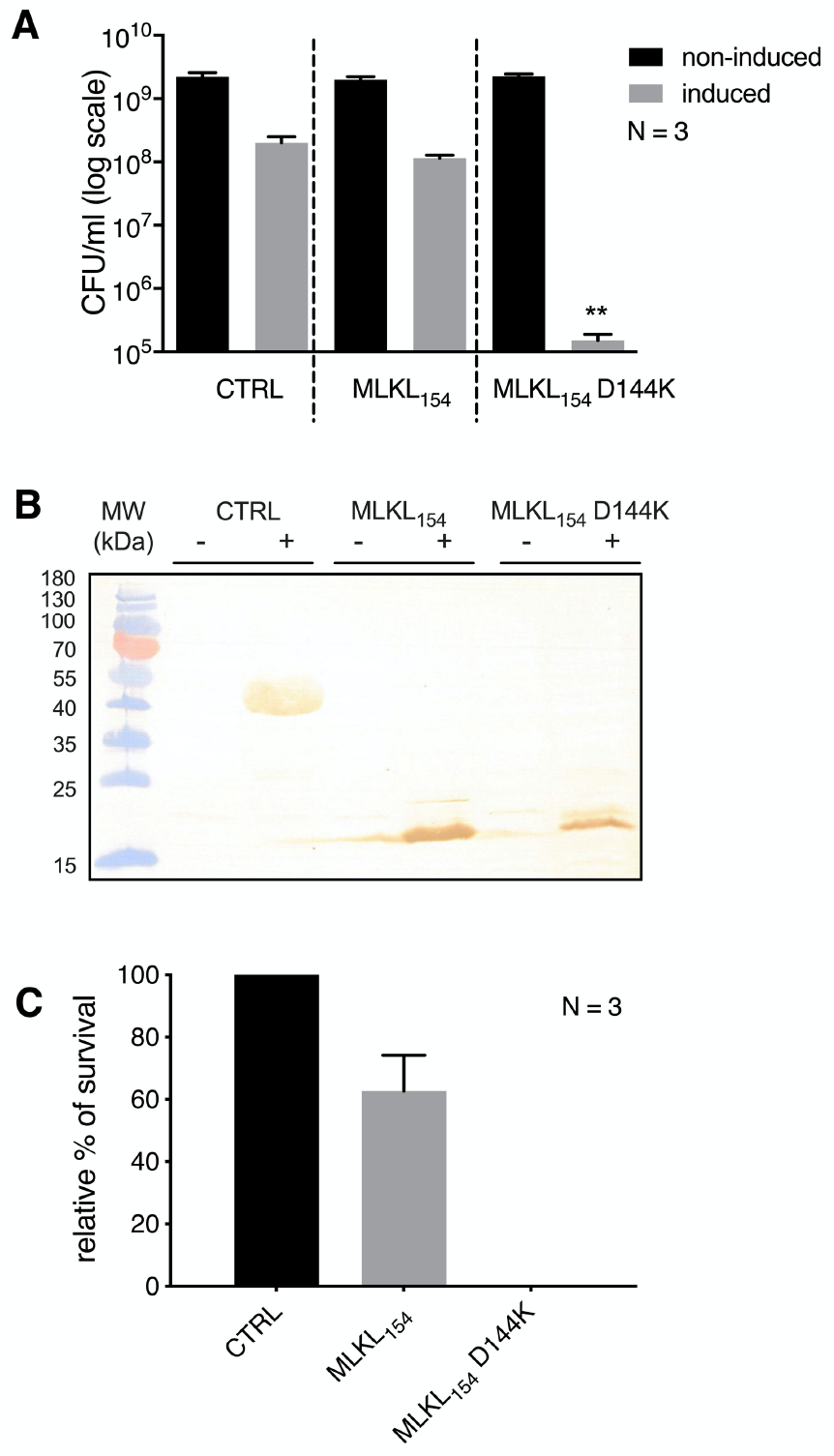
Over-expression of MLKL_154_ D144K lowers the viability of *E. coli*. (A) Viable CFU counts of *E. coli* BL21[DE3] pLysS transformed with an expression plasmid coding for the negative control, MLKL_154_, MLKL_154_ D144K were determined 2h post IPTG induction or non-induction. Shown is mean+S.E.M. of three independent experiments. (B) Protein samples of induced and non-induced bacterial were collected 2h after OD_600_ reached 0.55 and subjected to immunoblotting. (C) Relative bacterial survival was normalized to viable CFU count of induced control protein 2h post-induction. Shown is mean+S.E.M. of three independent experiments. Data are representative of three independent experiments. *P≤0.05, **P≤0.01.

Having established MLKL_154_ D144K lowers the viability of bacteria after its induction compared to MLKL_154_ and negative control, we decided to ectopically express MLKL_154_ and MLKL_154_ D144K in human embryonic kidney 239T cells (HEK293T). As we have previously shown, MLKL_154_ had no necroptotic effect on HEK293T survival. However, the necroptotic activity was activated when MLKL_154_ D144K was ectopically expressed (Fig. 3A) [33]. To see if the mutation D144K affects killing activity in the context of the full-length protein, we introduced the mutation into MLKL isoform 1 and performed concentration-dependent measurements. The necroptotic activity was always significantly higher for mutant compared to wild type MLKL1 protein (Fig. 3B). This strongly supports the notion that helix H6 is responsible for the inhibition of necroptotic activity. To further confirm this hypothesis, we ectopically overexpressed MLKL_139_, the N-terminal MLKL construct lacking the majority of the first brace helix H6 including D144, in HEK293T (Fig 3A). As predicted, removal of the majority of the first brace helix completely restored killing potential of the 4HB domain in HEK293T to the level of the mutant MLKL_154_ D144K (Fig. 3A).

**Figure 3:**
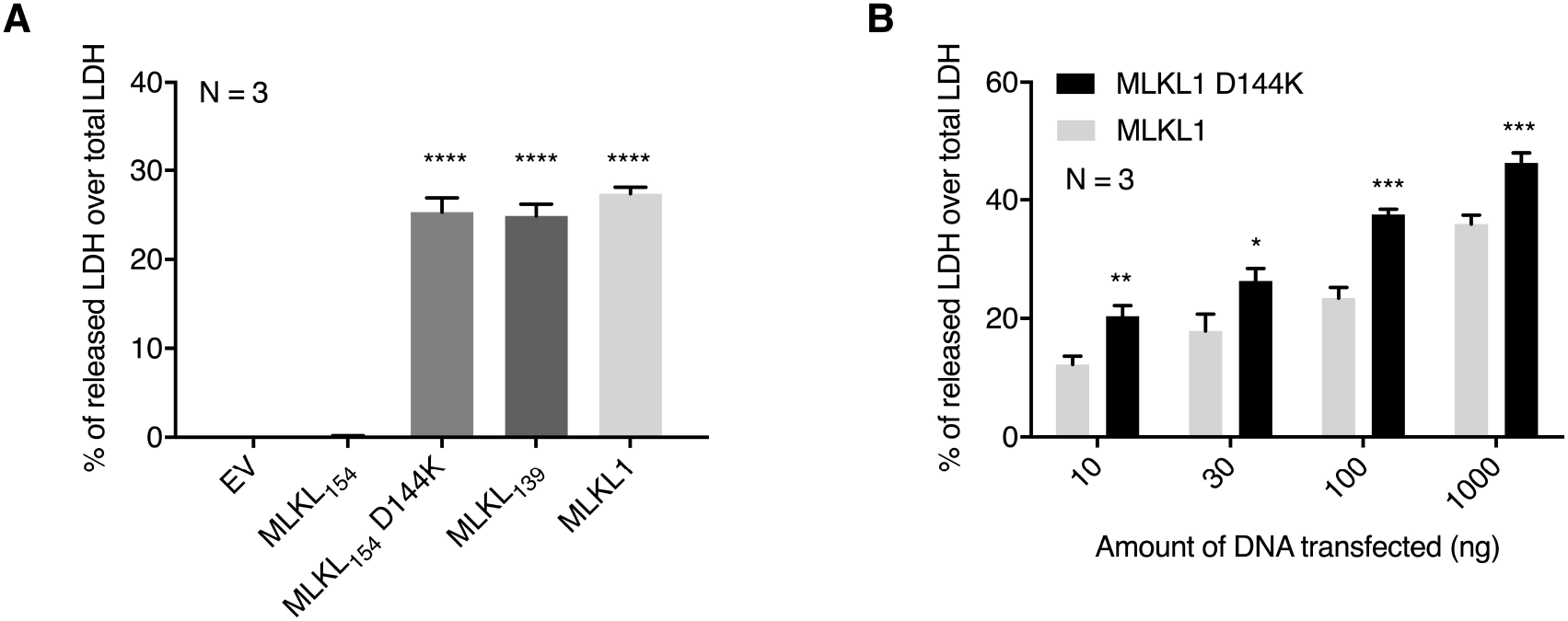
Single amino acid mutation D144K activates the necroptotic activity of the MLKL_154_ when ectopically expressed in HEK293T. (A) HEK293T were transfected with MLKL_154_, MLKL_154_ D144K, MLKL_139_ or EV (100ng). Cell viability was determined at 24h post-transfection by LDH release assay. (B) HEK293T were transfected with increasing amounts of expression plasmids encoding human MLKL1 and its mutant MLKL1 D144K with the total amount of transfected DNA being kept constant with EV. LDH release was measured at 24h post-transfection by LDH release assay. Shown is mean+S.E.M. of three independent experiments. *P≤0.05, **P≤0.01, ***P≤0.001, ****P≤0.0001.

### Mutation D144K disrupts packing interactions between helix H6 and 4HB domain resulting in decreased thermostability of MLKL_154_

It has been proposed that once 4HB domain binds PIPs, helix H6 becomes less structured [27]. To investigate whether the mutation of the conserved aspartate at position 144 affects interactions between helix H6 and 4HB domain, we measured the spectroscopic characteristics of MLKL_154_ and MLKL_154_ D144K. Purified recombinant proteins exhibited CD-spectra characteristic of α-helical proteins with minimum at 208nm and 222nm. This was in accordance with the NMR structure of the MLKL_2-154_ [26]. However, a substantial decrease in CD intensity was observed for MLKL_154_ D144K compared to MLKL_154_ (Fig. 4A), suggesting that helix H6 is partially unfolded. Indeed, deconvolution of CD-spectra revealed an α-helical content of approximately 63% (Table S1) for MLKL_154_ that decreased to 39% in the case of MLKL_154_ D144K. To further support these findings, we tested whether the addition of 2,2,2-trifluoroethanol (TFE) [40], an inducer of α-helix formation, reverts the effects observed by D144K mutation. Little or no difference was observed between CD-spectra in the presence or absence of 30% TFE for MLKL_154_ (Fig. 4A). However, the addition of 30% TFE increased CD intensity in MLKL_154_ D144K to a similar intensity observed for MLKL_154_.

**Figure 4:**
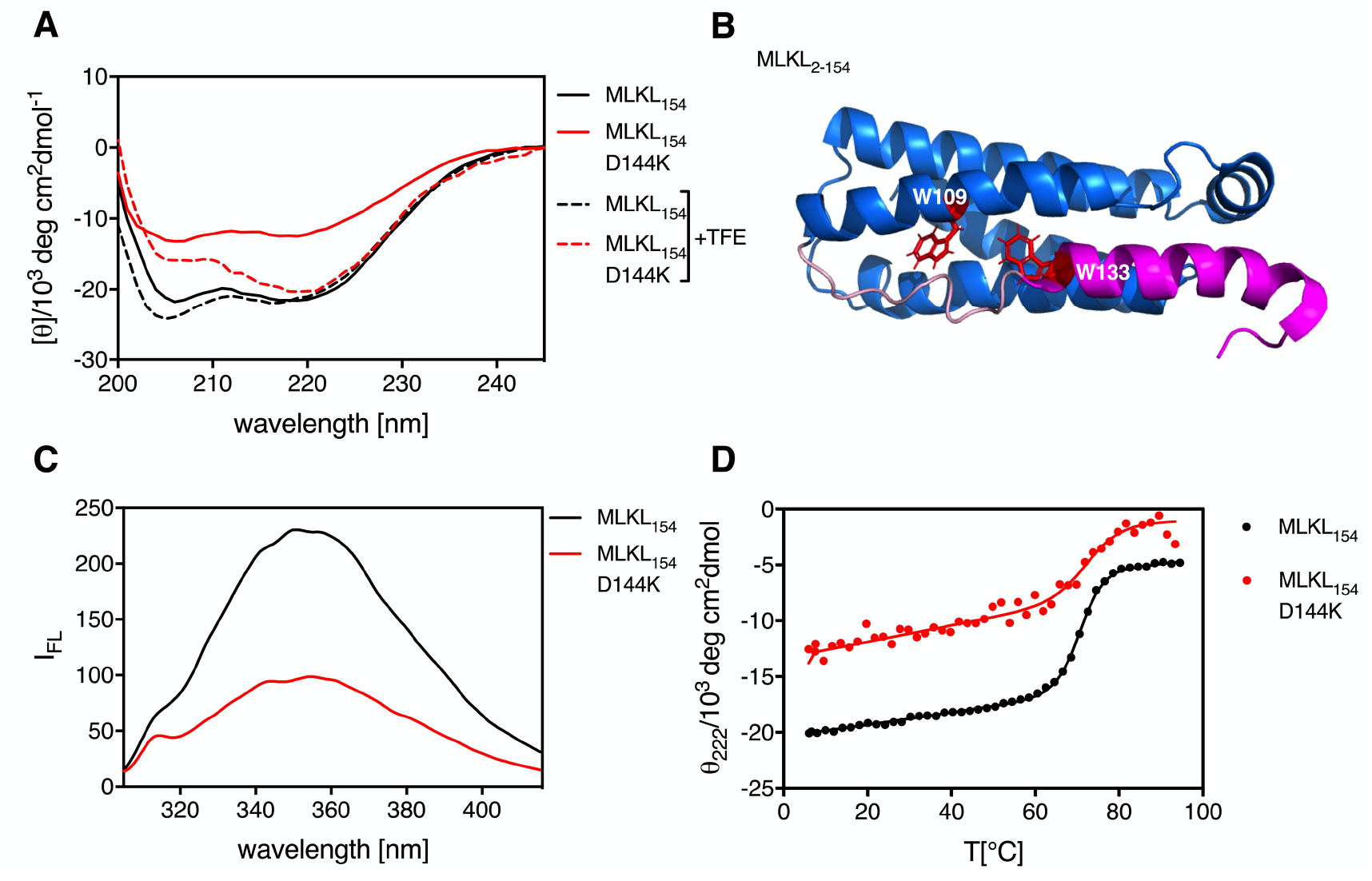
Single amino acid mutation D144K partially unwinds helix H6, disrupts its packing interactions with 4HB and destabilizes MLKL_154_. (A) Far UV CD-spectra of MLKL_154_ (black) and MLKL_154_ D144K (red) in 20mM acetate buffer pH 5.0 with 1mM DTT in the absence or the presence of 30% TFE (dashed lines). (B) Position of W residues in the structure of MLKL_2-154_ (PDB: 2MSV). W109 positioned within helix H5, and W133 positioned within helix H6 are presented in red sticks. Helix H6 of the brace region is shown in pink and 4HB is shown in blue. (C) Intrinsic fluorescence of MLKL_154_ (black line) and MLKL_154_ D144K (red line) at protein concentration 0.005mg/ml. (D) Thermal melts of MLKL_154_ D144K (red dots) and MLKL_154_ (black dots) recorded as CD intensity at 222nm in 20mM acetate buffer pH 5.0 with 1mM DTT. The two-state model analysis was performed (solid lines).

The MLKL_154_ has two tryptophan residues, one of which is located on helix H5, within 4HB domain and the other on helix H6 of the brace region (Fig. 4B). It is, therefore, reasonable to assume that conformational changes due to D144K mutation would be reflected in a change of intrinsic fluorescence which depends on solvent-exposure of tryptophan residues. Indeed, we observed decreased fluorescence intensity for MLKL_154_ D144K compared to MLKL_154_ (Fig. 4C). This indicates that the mutation D144K most likely disrupts packing interactions between helix H6 and 4HB domain, by partially exposing both W109 and W133 to solvent.

In light of these results, we investigated if the mutation has an effect on the thermodynamic stability of MLKL_154_. To address this question, we monitored changes in CD intensity as a function of temperature. According to the CD-spectra at 94°C, the majority of secondary structure for MLKL_154_ and MLKL_154_ D144K was lost in the denatured state. However, the secondary structure was only partially reversible for both proteins at pH 7.4 (Figure S1, 1A). Therefore, thermal scans were measured in acetate buffer at pH 5.0, which improved reversibility (Figure S1, 1B). This permitted us a reliable estimation of thermodynamic parameters using a two-state denaturation model (Fig. 4D). Stability of MLKL_154_ protein was estimated to 4.8kcal/mole at 25°C, and the introduced mutation D144K decreased protein stability to 3.7kcal/mol at the same temperature. The observed decrease in the stability was due to a lower enthalpic contribution for the D144K mutant. This was expected, since at least one hydrogen bond and favourable interactions of D144 with helix H2 were lost with D144K mutant. A part of decreased stability can also be attributed to the observation that helix H6 is partially disordered in the D144K mutant, given that helix unfolding is known to be accompanied by negative enthalpy change.

### Mutation of aspartate 144 enables MLKL_154_ D144K membrane translocation and possibly its oligomerization

Having shown the difference between the biological activity of MLKL_154_ and mutant MLKL_154_ D144K, we sought to determine whether there is a difference in subcellular localization between ectopically expressed wild type and mutant D144K protein in HEK293T. For this experiment, we fused both proteins with EGFP and analyzed their localization by confocal microscopy. As expected, MLKL_154_ was equally dispersed throughout cytoplasm 16h post-transfection. However, MLKL_154_ D144K has shown enrichment in cell membranes under the same conditions (Fig. 5A). Furthermore, mutant MLKL_154_ D144K formed discrete punctae 24h post-transfection, a phenomenon that was observed neither in cells transfected with MLKL_154_ nor in cells transfected with empty vector (Fig. 5B). Size exclusion chromatography (SEC) of purified tagless recombinant proteins revealed monomeric wild type and a mixture of monomeric and high molecular weight complexes (HMW) for D144K mutant protein. Similarly, SEC of tagless recombinant MLKL_139_ revealed a mixture of monomeric and high HMW (Figure S2).

**Figure 5:**
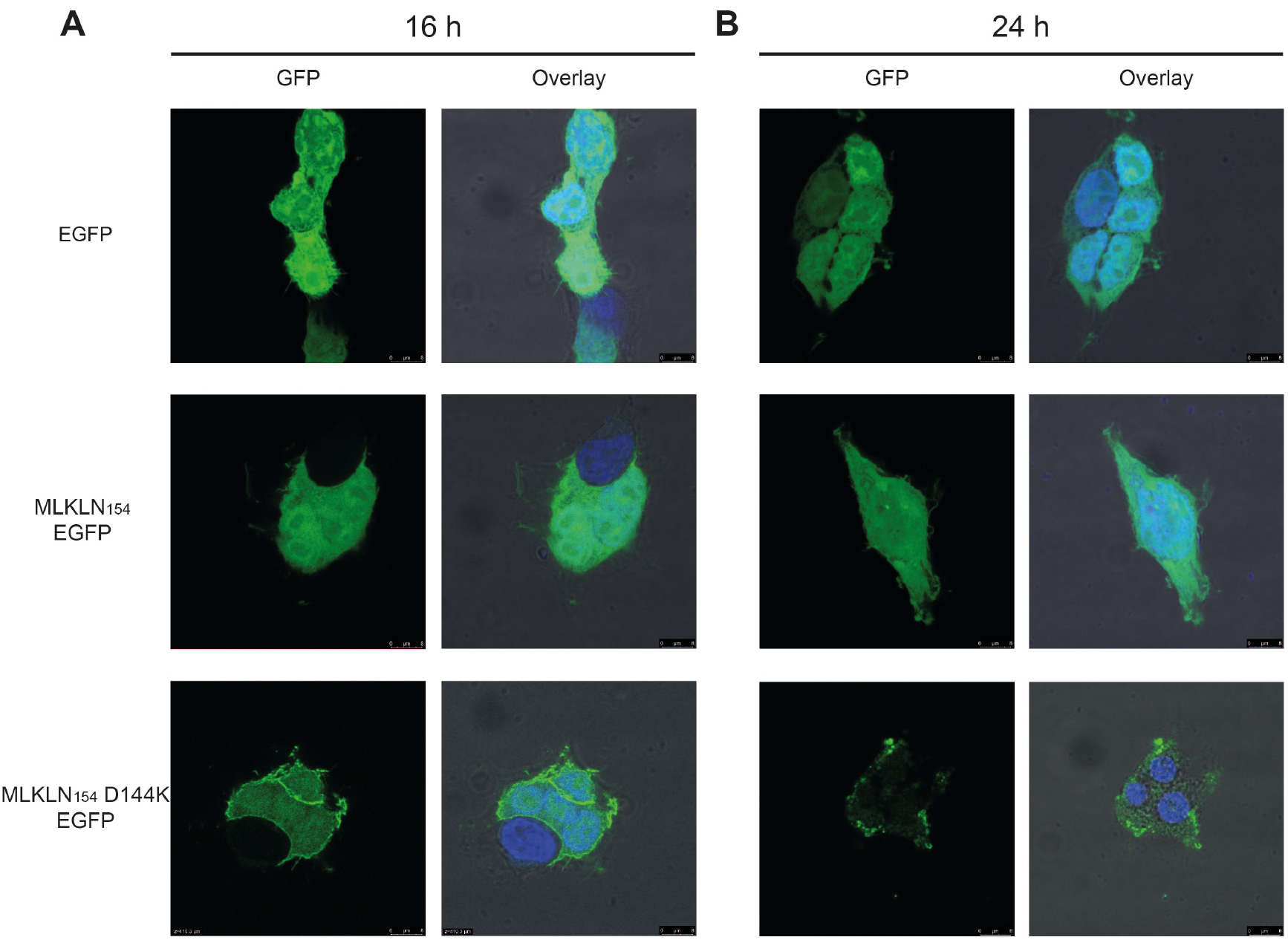
MLKL_154_ D144K is enriched in the plasma membrane and forms discrete punctae when overexpressed in HEK293T. HEK293T were transfected with 250ng of empty pEGFPN2 plasmid or pEGFPN2 coding for MLKL_154_ or MLKL_154_ D144K. EGFP and DAPI fluorescence were analyzed 16h (A) post-transfection or 24h (B) post-transfection. Similar results were apparent in three independent experiments.

## Discussion

MLKL consists of the N-terminal 4HB domain, the two-helix brace region and the C-terminal pseudokinase domain [19]. Even though both, the 4HB and the pseudokinase domain have been relatively well studied, little is known about the effect of the brace region on MLKL’s activation. Recently it has been reported that the second brace helix is involved in protein’s oligomerization [29], while the fist brace helix could have an inhibitory effect [26, 27, 33].

Previously we reported that the overexpression of the human MLKL 4HB domain with the intact brace region (residues 1-201) constitutively killed HEK293T cells [33]. Surprisingly, in the same study, we have shown that overexpression of truncated N-terminal MLKL (residues 1-154), a construct consisting of 4HB and the first α-helix of brace region completely compromised the ability to reconstitute cell death in HEK293T [33]. In order to elucidate how the first α-helix of brace region, helix H6 could regulate the activity of MLKL, we identified a well-conserved amino acid residue within this region, D144. We anticipated that a single amino acid mutation D144K would be sufficient to activate the killing potential of MLKL_154_ in HEK293T by moving helix H6 away from 4HB domain, possibly enabling protein’s oligomerization and membrane translocation.

The NMR structure of the human N-terminal domain (residues 2-154) revealed that the construct consists of the 4HB domain and the fist brace α-helix, helix H6 [26]. The results of liposome leakage assay showed significantly lower permeabilization activity of MLKL_2-154_ compared to MLKL_2-123_, a 4HB MLKL construct without the first brace helix [26]. This suggested a plug release mechanism, where helix H6 must be pulled away from 4HB domain to unleash its killing activity [26, 33]. Among 23 amino acid residues present in human MLKL, only aspartate 140 and 144 were conserved between 16 compared species. Single amino acid mutation D144K enabled inactive MLKL_154_ to become constitutively active when overexpressed in HEK293T. This was confirmed in the context of constitutively active full-length MLKL isoform 1 protein. Introduction of mutation D144K in MLKL1 resulted in the higher killing potential of mutant protein compared to wild type full-length MLKL. Similarly, missense mutation of mouse MLKL D139, amino acid residue equivalent to human D144, results in protein’s auto-activation [41].

Salt bridges between charged amino acids in helix H2 and helix H6 have previously been reported to have a role in plug release mechanism [20, 26]. Specifically, the double mutation of K26 and R30 (amino acid residues on helix H2 that could form salt bridges with helix H6) increased the liposome leakage compared to wild type MLKL_2-154_, indicating that salt bridges have a stabilizing effect on MLKL structure [26]. To further support this idea, mutations of R30 or E136 (amino acid residues forming a salt bridge between helix H2 and helix H6) induced necroptosis better than wild type full-length MLKL [27]. Amino acid residue D144 is positioned at the C-terminal end of the helix H6. This region interacts with the N-terminal region of helix H2, composed of C24, K25 and K26. D144 is within the hydrogen bond distance to C24, and D144 side chain is positioned precisely over the top of the helix H2. Helix H2 dipole and its two lysine residues (K25 and K26), although not involved in salt bridges with D144, strengthen this interaction. Our data from tryptophan intrinsic fluorescence support the idea that the mutation D144K facilitates helix H6 to be moved away from the 4HB domain. Furthermore, the mutation D144K decreased α-helical content of the protein by 23%. Addition of TFE, a well know secondary structure stabilizer, partially reversed the effects induced by D144K mutation, since helicity became comparable to those observed for the wild type protein. Estimation of thermodynamic parameters revealed lower thermodynamic stability of MLKL_154_ D144K compared to MLKL_154_. Together, these results show that the introduced single amino acid mutation promotes helix H6 being moved away from 4HB domain and possibly induces intrinsically disorder of helix H6. Indeed, Quarato et al. (2016) proposed a mechanism of MLKL membrane binding that induces brace helix H6 displacement and its disorder. In their model, a putative high-affinity site for PIP binding on helix H2 is masked by helix H6. Once MLKL binds to the plasma membrane via a low-affinity PIP binding site on helix H1, MLKL undergoes structural rearrangement where helix H6 is moved away from helix H2, and high affinity PIP binding site on helix H2 is unveiled. Their NMR TROSY analysis of MLKL_156_ in the presence of PIPs showed that upon lipid binding helix H6 becomes unstructured. These results are well in line with our spectroscopic data. Moreover, we show overexpression of MLKL_139_, a construct lacking a part of helix H6 that would hide the possible putative PIP binding site on helix H2, is constitutively active in HEK293T.

MLKL’s plasma membrane translocation is a hallmark of necroptosis [21–25, 42, 43]. The N-terminal domain of MLKL is responsible for membrane phosphatidylinositol phosphate and cardiolipin binding [22, 25]. Dondelinger et al. (2014) showed EGFP tagged full-length human MLKL and MLKL_1-180_ were translocated to plasma membranes when overexpressed in HEK293T. This translocation was not observed for EGFP tagged C-terminal MLKL domain (amino acid residues 181-471), the construct that had no constitutive killing activity in HEK293T. Comparably, our results showed no plasma membrane translocation for EGFP tagged MLKL_154_, the inactive construct, 16 or 24h post-transfection in HEK293T. MLKL_154_ was similarly distributed in the cytoplasm as negative control, empty vector constitutively expressing EGFP. However, plasma membrane translocation was observed in case of EGFP tagged mutant MLKL_154_ D144K 16h post-transfection in HEK293T. More importantly, 24h post-transfection mutant MLKL_154_ D144K formed discrete foci in the membrane region. This could indicate that the mutant protein formed oligomers. Indeed, N-terminal domain has been reported to play a role in MLKL’s oligomerization. Human recombinant 4HB (amino acid residues 1-140) and 4HB with first brace helix (amino acid residues 2-154) do not form oligomers [26, 27]. Same properties were observed for mouse N-terminal domain (first 158 amino acid residues). However, the addition of the second brace helix resulted in homotrimer formation *in vitro* [29]. Quarato et al. (2016) showed that a full brace region (residues 141-182) is responsible for oligomerization. These results suggest that second brace helix enables oligomerization, and the first brace helix prevents it. Recombinant tagless human MLKL_154_ is monomeric as judged by the size-exclusion chromatography. However, MLKL_154_ D144K exists as a mixture of monomers and high molecular weight complexes. Same polydispersity has been observed for MLKL_139_, construct lacking a majority of first brace helix. These results support the hypothesis that first brace helix prevents oligomerization, while 4HB domain alone might be somehow important for oligomerization. This hypothesis is supported by experiments from Dondelinger et al. (2014). Overexpression of constitutively active 4HB (amino acid residues 1-125) in HEK293T formed HMW complexes. HMW complexes were observed in the case of all constitutively active MLKL constructs including MLKL_1-125_, MLKL_1-180_, MLKL_1-210_ and full-length MLKL. Importantly, HMW constructs were not observed in the case of inactive MLKL_1-167_ [22]. In the present work, we identified amino acid residue responsible for the structural stabilization of the N-terminal domain of MLKL. By mutating this amino acid, we confirmed the hypothesis of helix H6 constraining necroptotic activity. Collectively, our results demonstrate that brace helix does not only have an interdomain linker role, but it is actively involved in the regulation of MLKL’s activation.

## Author Contributions

KHS, GG, SH, JL and MPK performed or assisted with experimentation. KHS, SH and MPK were involved in experimental analysis and interpretation. GG, KHS, SH and JL contributed to the writing of this manuscript.

## Abbreviations

MLKL: Mixed lineage kinase domain-like
RIP: Receptor-interacting protein kinase
MLKL1: MLKL isoform 1
MLKL154: MLKL N-terminal domain of 154 amino acid residues
MLKL139: MLKL N-terminal domain of 139 amino acid residues
MLKL154 D144K: MLKL N-terminal domain of 154 amino acid residues with mutation D144K
EV: empty vector
4HB: four-helix bundle domain
helix H6: α6 helix
aa: amino acid
PIP: phosphatidylinositol phosphate
EGFP: enhanced green fluorescent protein
CD: Circular dichroism spectroscopy

