## Supplemental Figure1, Supplemental Figure 2, Supplemental Table 1 for "MLKL D144K mutation activates the necroptotic activity of the N-terminal MLKL domain"

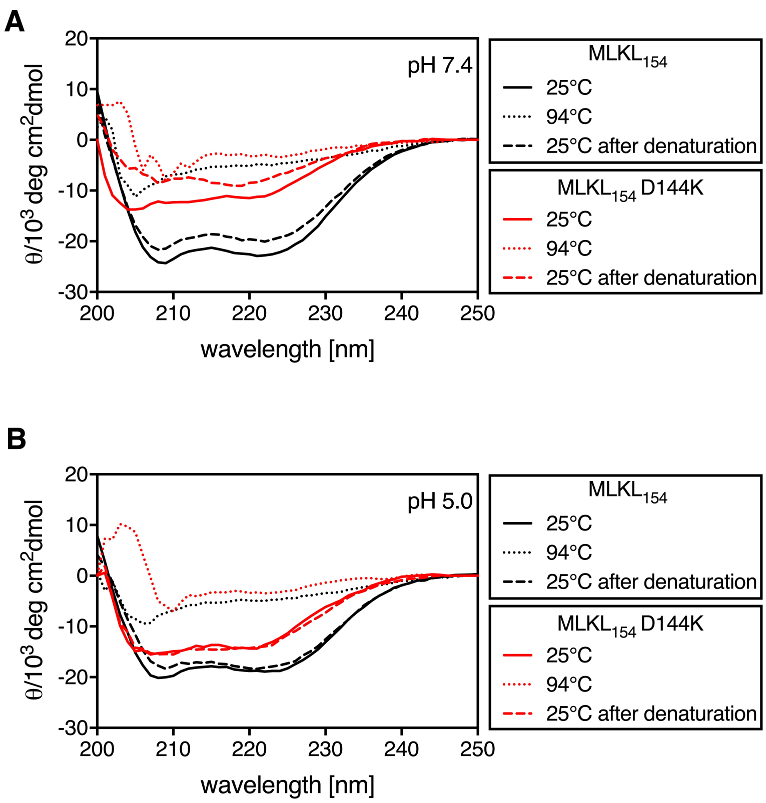


**Figure S1: Reversibility of secondary structure for MLKL$_{154}$ and MLKL$_{154}$ D144K proteins is improved at pH 5.0 compared to pH 7.4.** CD-spectra of MLKL$_{154}$ and MLKL$_{154}$ D144K were measured at 25°C, 94°C and 25°C after denaturation at pH 7.4 (A) and pH 5.0 (B) with 1mM DTT. All samples were measured at 0.2mg/ml for MLKL$_{154}$ and 0.01mg/ml for MLKL$_{154}$ D144K.

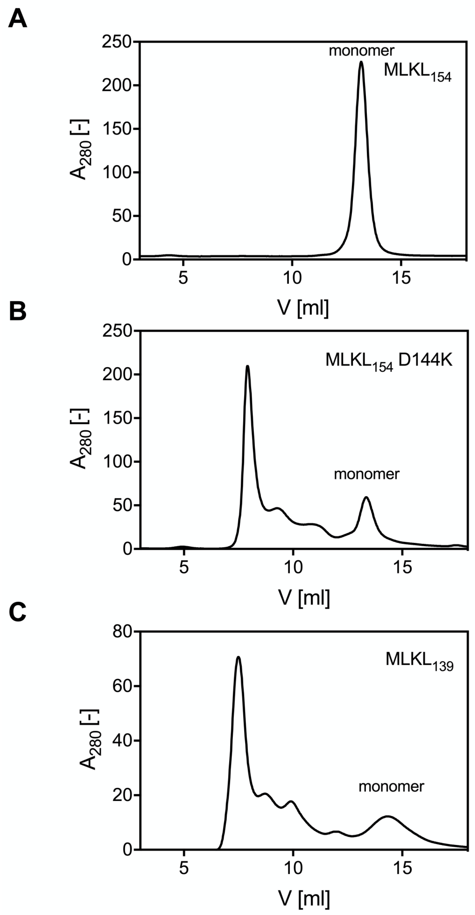


**Figure S2: Size exclusion chromatography of $MLKL_{154}$, $MLKL_{154}$ D144K and $MLKL_{139}$**. All samples were measured in 50mM Tris-HCl pH 7.4, 150mM NaCl and 1.5mM DTT. The total amount of loaded purified recombinant $MLKL_{154}$ and $MLKL_{154}$ D144K was 1.3mg and 0.8mg for $MLKL_{139}$.

**Table S1: Content of CD-spectra for MLKL154 and MLKL154 D144K.** CD-spectra of MLKL154 or $MLKL_{154}$ D144K was calculated using BeStSEL at pH 7.4. and pH 5.0 under reducing (1mM DTT) or non-reducing conditions.

| | **$MLKL_{154}$** | | | | **$MLKL_{154}$ D144K** | | | |
|---|---|---|---|---|---|---|---|---|
| | Reducing conditions | | Non reducing conditions | | Reducing conditions | | Non reducing conditions | |
| | pH 7.4 | pH 5.0 | pH 7.4 | pH 5.0 | pH 7.4 | pH 5.0 | pH 7.4 | pH 5.0 |
| Helix [%] | 74.8 | 63.0 | 58.6 | 60.8 | 47.4 | 39.2 | 37.4 | 43.5 |
| Antiparallel [%] | 0.0 | 13.9 | 3.8 | 19.6 | 11.5 | 44.4 | 24.7 | 5.4 |
| Parallel [%] | 0.0 | 0.0 | 0.0 | 0.0 | 0.0 | 0.0 | 0.0 | 4.3 |
| Turns [%] | 14.5 | 13.3 | 10.8 | 11.2 | 19.4 | 16.4 | 16.3 | 9.4 |
| Other [%] | 10.8 | 9.8 | 26.8 | 8.3 | 21.7 | 0.0 | 21.6 | 37.3 |
